# MitoZ: A toolkit for mitochondrial genome assembly, annotation and visualization

**DOI:** 10.1101/489955

**Authors:** Guanliang Meng, Yiyuan Li, Chentao Yang, Shanlin Liu

## Abstract

Mitochondrial genome (mitogenome) plays important roles in evolutionary and ecological studies. It becomes routine to utilize multiple genes on mitogenome or the entire mitogenomes to investigate phylogeny and biodiversity of focal groups with the onset of High Throughput Sequencing technologies. We developed a mitogenome toolkit MitoZ, consisting of independent modules of *de novo* assembly, findMitoScaf, annotation and visualization, that can generate mitogenome assembly together with annotation and visualization results from HTS raw reads. We evaluated its performance using a total of 50 samples of which mitogenomes are publicly available. The results showed that MitoZ can recover more full-length mitogenomes with higher accuracy compared to the other available mitogenome assemblers. Overall, MitoZ provides a one-click solution to construct the annotated mitogenome from HTS raw data and will facilitate large scale ecological and evolutionary studies. MitoZ is free open source software distributed under GPLv3 license and available at https://github.com/linzhi2013/MitoZ.

## INTRODUCTION

With the onset of High Throughput Sequencing (HTS) technologies, we have entered an era in which massive nucleic acid sequencing is becoming routine in phylogenetic and biodiversity monitoring studies (1,2). For example, metabarcoding studies, by taking advantage of complex DNA extracts (e.g. environmental DNA (eDNA)), identify multiple taxa simultaneously from diverse types of samples – stomach contents (3), feces (4,5), sediments (6), soil or water (6-8). In most cases, these studies deal with degraded DNA, thus are in urgent demand for short barcoding fragments for taxonomic identification (9,10). Genes on mitochondrial genomes are preferred due to high copy number per cell, making them more likely be picked up than single-copy nuclear genes. Rapid access to mitochondrial genomes of a myriad of taxa will, firstly, provide critical taxonomic connections between the most abundant and well-constructed DNA barcode COI and those eDNA widely-adopted short markers *12S rRNA, 16S rRNA, CYTB* et al. (reviewed by 11); secondly, facilitate the fast-emerging approach – mito-genomics (12-14), which circumvents PCR and requires a taxonomically well covered reference dataset used both for species identification and in gene capture array design (2,15). In addition to its importance in biodiversity monitoring, mitochondrial genome also records maternal inheritance information and is extensively utilized to infer phylogenetic relationship between diverse lineages (1,16).

Apart from the mitogenomes achieved using long-range PCR followed by primer walking strategy and Sanger dideoxy sequencing (17), quite some mitogenomes were obtained using a reference-based method via HTS platform (e.g. 18,19-21). Traditional genome assembly software, for instance, SOAPdenovo2 (22), ALLPATHS-LG (23), Platanus (24), can hardly assemble complete mitogenomes since they are programmed to abandon sequences with extremely-high depth. The two frequently-used mitogenome assembly software, MITObim (25) and NOVOPlasty (26), require closely-related mitochondrial fragments as seeds to anchor short reads and build initial datasets. However, it is often difficult to set an appropriate criterion to define closely-related species – e.g. should an appropriate criterion be congeneric or coordinal in the Linnaean system. The similarities between species also vary a lot between different groups (27). There are also some genera within which none species has a complete mitogenome albeit the plunging cost of sequencing (28). In addition, both software can only generate mitogenome assembly as their final outputs. Thus, separate software, like DOGMA (29), MOSAS (30), MITOS (31) are required for the following genome annotation (32). Besides, all of the three aforementioned annotation software only provide web page version and can hardly deal with assembly with multiple scaffolds.

Here we presented a mitochondrial genome toolkit, MitoZ, providing a one-click solution from HTS raw reads to genome assembly together with annotation and visualization outputs. MitoZ is programmed in Python3 (33) with the assembly module of a modified version of SOAPdenovo-Trans (34), the annotation module of a PERL based script for protein coding genes (PCGs), MiTFi (35) for tRNA and infernal-1.1.1 (36) for rRNA (Fig. 1). We tested the accuracy and efficiency of MitoZ using a batch of mammals and arthropods which have both mitogenomes obtained by Sanger sequencing in NCBI RefSeq database (37) and shotgun Paired-End reads in NCBI Sequence Read Archive (SRA) database (38). The results showed that MitoZ can recover 97.33% of PCGs and rRNA genes of the test samples, of which 94.66% genes are in full length and the recovered genes are of high similarity (≥97%) to their sanger sequenced mitogenome.

**Fig. 1.**
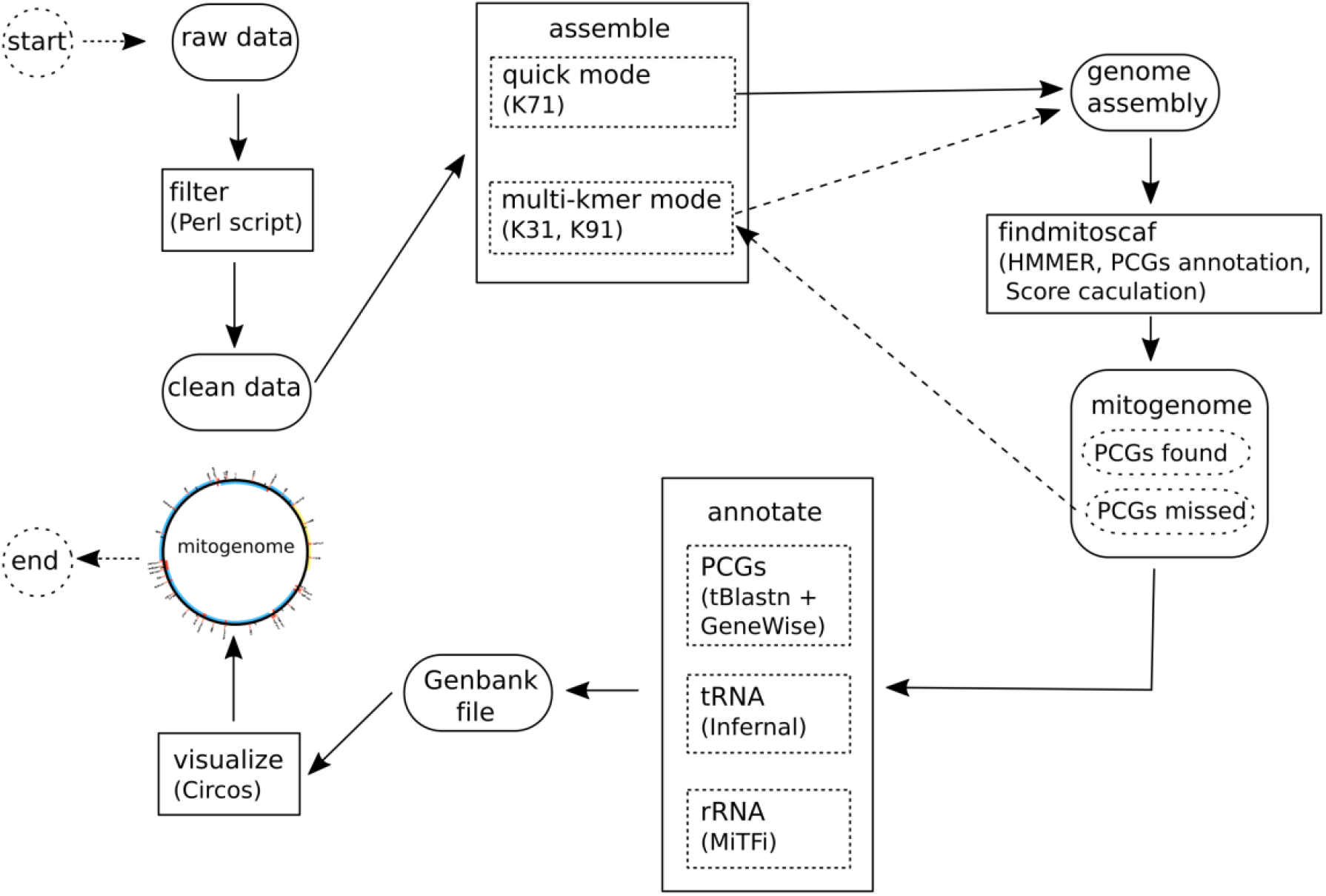
MitoZ toolkit components. Ellipses indicate input data files, rectangles (solid line) represent functional modules in MitoZ which can be run independently when users provide corresponding input files.

## MATERIAL AND METHODS

### 1 Samples for test

A total of 30 arthropod and 26 mammal species (Table S1) were selected to evaluate the performance of MitoZ. Species were picked up by considerations: 1) have mitogenome in NCBI RefSeq database (37) and were obtained using traditional Sanger sequencing method (refer as sanger mitogenomes afterward); 2) have HTS data with a volume size ≥ 3 Gb and Paired-End (PE) read length ≥ 91 bp in NCBI Sequence Read Archive (SRA) database (38); 3) For mammal, data generated from tissue samples were preferred, but 7 blood samples were included as well for comparison.

We estimated the ratio of mitochondrial derived reads (MDR) for each sample by aligning raw reads to their corresponding sanger mitogenomes using BWA (version 0.7.12-r1039) (39). It showed that most samples had a MDR ratio in a range from 0.12% to 0.51% with the mammal blood samples possessing a significant lower MDR ratio from 0.01% to 0.05% (Table S2). In addition, we also noticed that six samples (including three mammal non-blood samples, two mammal blood samples and one arthropod sample) possessed MDR ratio of zero. For those samples, the MDR could be removed on purpose before data deposit. Thus, we removed them from the following performance evaluation, leading to a total of 50 samples in the final dataset, consisting of 29 arthropods, 16 mammal non-blood samples and five mammal blood samples. See Table S3 and Table S4 for details of the procedures of species selection and dataset download.

### 2 MitoZ

MitoZ consists of multiple modules, including raw data pretreatment, *de novo* assembly, candidate mitochondrial sequences searching, mitogenome annotation and visualization (Fig. 1). Each module can work independently in case users want a sub part of the entire workflow.

#### 2.1 Raw data pretreatment

A Perl script is designed for filtering the raw data generated by HTS platforms, such as Hiseq 4000. It accepts Pair-End (PE) reads and filters out reads with many ‘N’s, low qualitie reads, or reads of PCR duplicates (defined as a pair of identical reads). By default, reads will be removed if: 1) of more than 40% low quality (Q<=17) bases; 2) of more than 10 Ns; 3) are PCR duplications

#### 2.2 *De novo* assembly

Algorithms adopted in SOAPdenovo-Trans (34) fit well for mitogenome assemblies from nuclear DNA extracts, where mitogenome sequences possess considerable higher copy number comparing to nuclear genome sequences - being alike to the differential gene expression patterns it is designed for. We adopted all the algorithms in SOAPdenovo-Trans except for the scaffolding step where we ask for higher linkage support to avoid connections between mitochondrial reads and nuclear mitochondrial DNA segments (NUMTs) (40) and to improve accuracy.

MitoZ has two assembly modes – Quick mode and Multi-Kmer mode. It uses the Quick mode by default, where only one kmer size (K=71) is used for assembly. Users can also use the Multi-Kmer mode to search for the missing PCGs (if any) failed in the Quick mode.

#### 2.3 Mitogenome sequences identification

The assemblies contain both mitogenome sequences and nuclear genome sequences. Thus, we filter out candidate mitogenome sequences using a profile Hidden Markov Model (profile HMM) (41,42) based method which works in an efficient and criteria-relaxed way at first. HMMER (version 3.1b2) (http://hmmer.org/) (43) is utilized to construct HMM models for both mammal and arthropod clades (2413 and 4007 species, respectively, see Table S5) in the current version. Then, we conduct PCG annotation for candidate mitogenome sequences. The PCG annotation is detailed in section 2.2.4. After that, we use the following three steps to remove potential false positive mitochondrial scaffolds, such as NUMTs and contaminations:

1. Each candidate mitogenome sequence was assigned to a Linnaean taxonomic name using a Python package ETE3 (44) according to their most closely-related PCG homologs in the PCG annotation step. Then, filter out sequences falling outside of a user predefined taxonomic rank, which can be set as order, family or genus.
2. We calculated a confident score *S*_*j*_ for each scaffold to determine the final mitogenome sequences. *S*_*j*_ is calculated using a formula as follows:

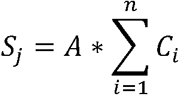

where *A* represents the assemble reliability of each assembly. It is a weight factor representing the reliability of *de bruijin* route selection – the higher the better. Its value is mainly consisted of two factors - Kmer depth of contigs and read supportive number when connect contigs to scaffolds. For sequences generated using other assembly software, *A* will be surrogated by average depth information calculated according to the number of reads that can be mapped to the targeted sequences, *n* indicates the number of PCGs in sequence *j* and C_*i*_-indicates the completeness (in percentage) of *i* th PCG, which is calculated by dividing the length for each gene by the length of its shortest reference counterpart in the annotation database.
3. Sequences are ranked by their confident scores. Then, MitoZ tries to find all 13 PCGs from the sequence with the highest confident score. In case mitogenome is not assembled as an integrate single sequence, MitoZ finds the rest PCGs from sequences by rank and skips sequences containing conflict genes, e.g. a complete *COX1* gene is located in a former sequence, then the latter sequences with lower *S*_*j*_ score containing *COX1* gene (complete or not) will be skipped. However, if former *COX1* is incomplete, the latter sequences containing also an incomplete *COX1* genes will be kept for PCG searching. The searching stops when 13 PCGs are all located, or sequences are run out. In addition, Sequences, regardless of gene conflicts, containing ≥ 5 PCGs (complete or not) will be retained to confirm identities, e.g. parasites.

#### 2.4 Genome annotation

##### protein coding genes

An in-house Perl script is designed for PCG annotation. Basically, the script finds candidate PCG sequences by aligning nucleotide sequences to a local protein sequences database using tBlastn in BLAST (version 2.2.19) (45), then uses *Genewise* (version 2.2.0) (46) to determine the boundaries of each PCG. MitoZ further tries to determine the precise position of start codons and stop codons by translating the nucleotide sequence with proper mitochondrial genetic code. MitoZ tries to find ‘TA’ or ‘T’ bases, assuming TAA stop codon is completed by adding 3’ A residues to the mRNA in case of absence of the standard stop codons. The current version includes protein database of both Mammals and Arthropods (Table S6).

##### tRNA genes

MiTFi (35), a covariance models (CMs) (42) based method, is adopted to annotate mitochondrial tRNA genes. It contains 22 manually curated tRNAs database for metazoan and employs a step-wise procedure to search for tRNA candidates in query sequences. MitoZ, by default, outputs tRNA annotation results of e-value ≤ 0.001 by setting MiTFi parameters as “-cores 1 -evalue 0.001 -onlycutoff -code 2/5(representing Mammal/Arthropod)”

##### rRNA genes

The 12S *rRNA* and 16S *rRNA* genes are annotated using infernal-1.1.1 (36) with the published rRNA CM models (31). MitoZ searches for rRNA with the local searching mode implemented in infernal-1.1.1 in case no candidates are detected using the global searching mode.

#### 2.5 Visualization

Mitochondrion genome features can be illustrated with an independent module, which employed Circos (47) to depict gene elements features, such as PCGs, rRNA genes, tRNA genes, GC content, and sequencing depth distribution. The color of each element can be set as personal preference.

### 3 Performance Evaluation

#### MitoZ assembly

We firstly applied MitoZ quick mode to all the 50 test samples, and tried to filter out contamination sequences by mitochondrial PCG annotation (set --requiring_taxa to be taxonomic rank of order for each species). For those didn’t get 13 PCGs, we applied another run using multi-Kmer mode.

#### Assembly quality

We examined the similarities between those mitochondrial genes (PCGs + rRNA) MitoZ recovered and that of Sanger mitogenomes using megablast in BLAST+ (2.6.0) (48). The genes with similarities < 97% to their sanger counterparts were regarded as false positive, but those false positives who can find matches (similarity≥ 97%) to their corresponding genes of the same species on NCBI (detailed in Table S7 and Table S8 and Table S9) were regarded as correct assemblies in the following statistics. We used MAFFT (version 7.309) (49,50) and Unipro UGENE (version 1.26.1) (51) to conduct global alignment between each pair and check mismatches, respectively.

#### Factors influencing mitogenome assembly

A+T content plays an important role in HTS experiments since the known bias in the library preparation step leading to genome regions that possess extreme A+T content tend to have low sequencing depth and are difficult to be assembled (52). Plus, the heterozygosity rate works against the genome assembly quality using HTS platforms (53). The “heteroplasmy” of our samples may come from pooled individuals aiming to produce enough DNA extracts for HTS library construction. We investigated the influences of sequence characteristics on the assembly qualities, including MDR ratio, depth, A+T content and mitogenome heteroplasmy. We aligned HTS reads on their corresponding mitogenomes using BWA (39) and calculated regional depth using SAMtools (54). We calculated the heterozygosity value for each sample based on Site Frequency Spectrum (SFS) obtained using ANGSD (55).

#### Comparison between MitoZ and NOVOPlasty

We also ran NOVOPlasty (version 2.7.2) to assemble mitogenomes of the 50 test samples with default parameters and the corresponding sanger mitogenome of each species was used as the reference seed. We then annotated the NOVOPlasty results with the annotate module in MitoZ and examined the similarities between those mitochondrial genes (PCGs + rRNA) obtained by NOVOPlasty and that of Sanger mitogenomes (detailed in Table S10 and Table S11).

## RESULTS

### Mitogenome completeness

Of the 750 genes ((13 PCGs + 2 rRNAs) * 50 species), 691 (92.13%) genes were full-length recovered, 39 genes (5.20%) were partially recovered and 20 (2.67%) genes were not assembled by MitoZ (Fig. 2 (iii)). The Multi-Kmer mode contributed a total of 46 genes that were either failed (33) or partially (13) assembled in the Quick mode.

**Fig. 2.**
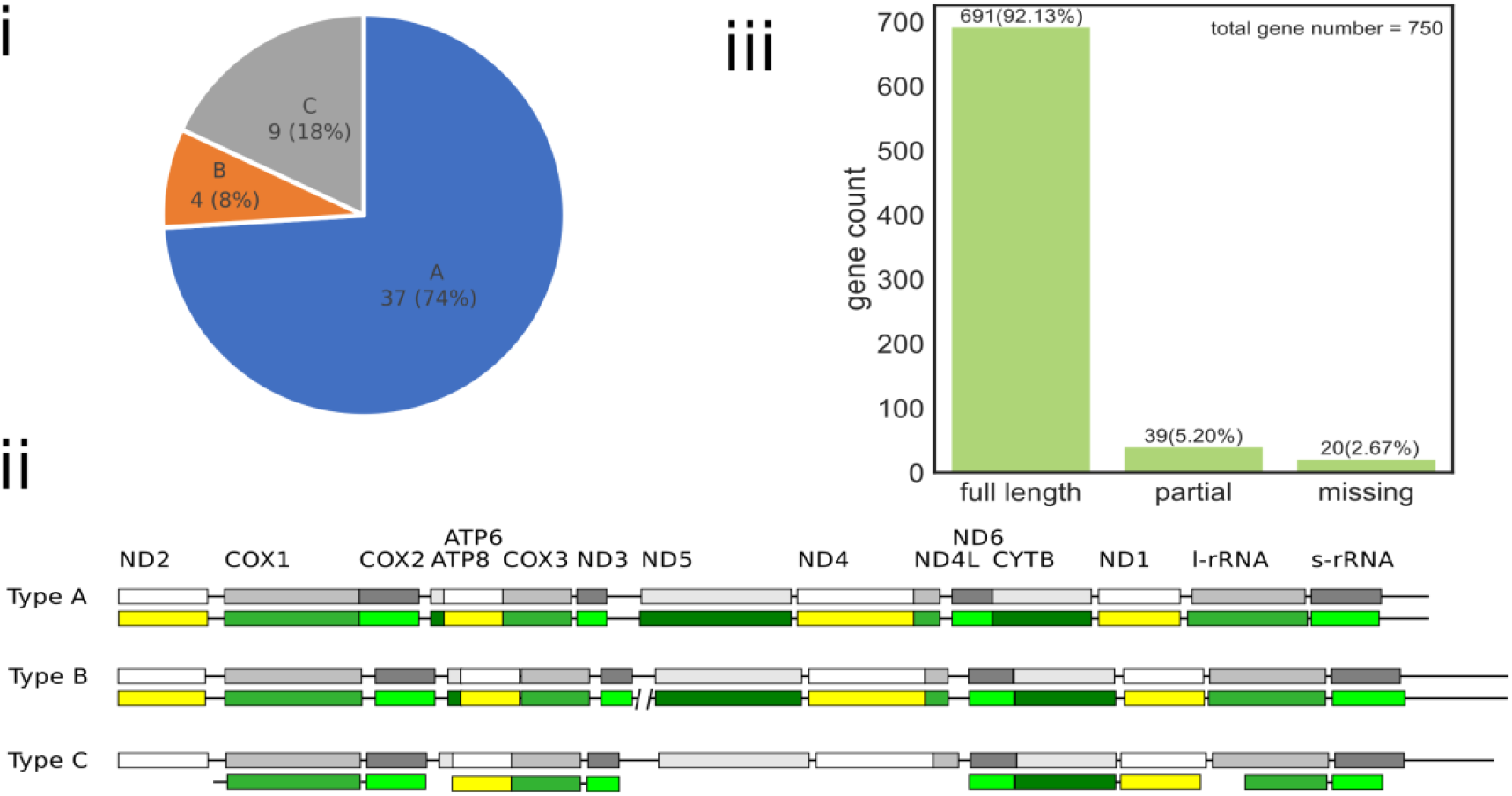
MitoZ test result. (i) the proportion of assembled mitogenomes in different assembly categories; (ii) diagram of different assembly types where the boxes represent PCGs or rRNA genes and the solid lines stand for the other parts of mitogenomes and the upper grey boxes represent sanger mitogenome while the lower colorful ones are assembled by MitoZ; (iii): the gene completeness distribution of PCGs and rRNA genes of the 50 test species. “full length” means gene completeness ≥ 95%, “partial” means gene completeness < 95%, and “missing” means the genes that were not recovered by MitoZ.

We categorized our assembly results into four types: 1) Type A, all the genes (13 PCGs and 2 rRNA) were recovered and represented by one single sequence; 2) Type B, all the genes were assembled, however represented by ≥ 2 sequences; 3) Type C, not all but more than half (≥ 8) of the genes were recovered; 4) Type D, the number of recovered genes was less than eight. In total, we got 37 (74.00%) mitogenomes of type A, four (8.00%) mitogenomes of type B, nine (18.00%) mitogenomes of type C, and none mitogenome of type D. See Fig. 2 (i) and (ii).

### Gene similarities

For the 735 PCGs and rRNA genes recovered, 724 (98.50%) genes matched their Sanger counterparts well (similarity≥97%, Fig. 3). We further checked these single nucleotide variances (SNV) between genes assembled by MitoZ and their Sanger counterparts. Although the SNVs can arise from individual variances or mitochondrial heteroplasmy, it is also worth to note that those non-perfect-match genes always possess high sequencing depth in HTS assemblies except for the ones located around “Ns” regions (Fig. S1 (i)) and those SNVs are in most cases located in homopolymers or A+T-rich regions (Fig. S1 (ii)), where are regions that typical Sanger sequencing errors happen.

**Fig. 3.**
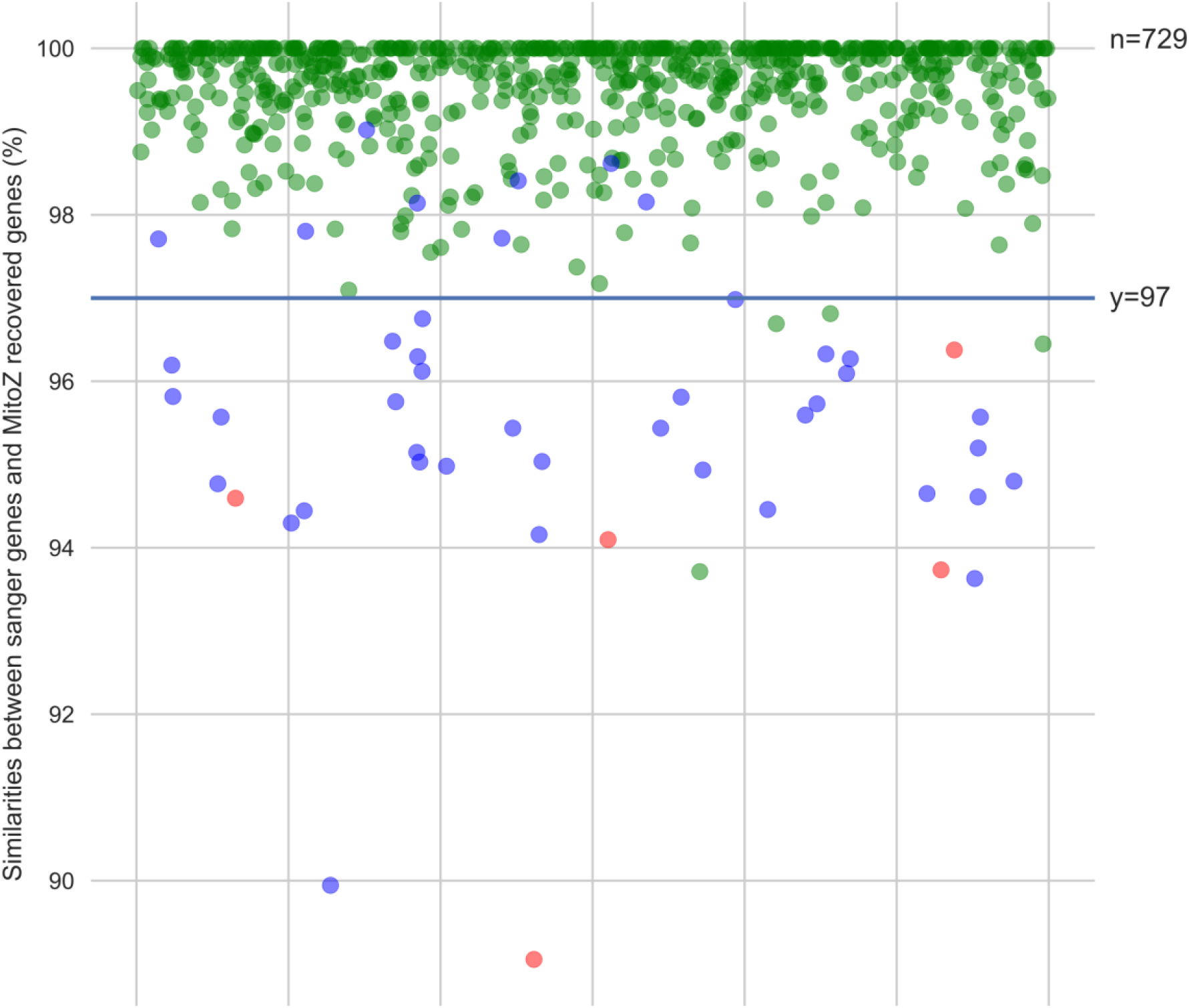
Gene similarities (MitoZ vs Sanger). The blue dots present three species whose genes possessed similarities < 97% to their Sanger mitogenomes but can find high similarity genes of the same species in NCBI NT database. Such incongruences could derive from intraspecies variances. The red points (5 in total) present genes possessed similarities < 97% to their Sanger mitogenomes and could not find better hits in NCBI. The rest genes and samples were presented by green dots.

The 5 false positive genes (the red dots in Fig. 3) with low similarities to their Sanger counterparts could be contributed to insufficient sequencing depth in HTS sequencing and sequencing errors in Sanger mitogenomes, see Table S9 for details. Fig. 4 shows an example of mitogenome visualization by MitoZ.

**Fig. 4.**
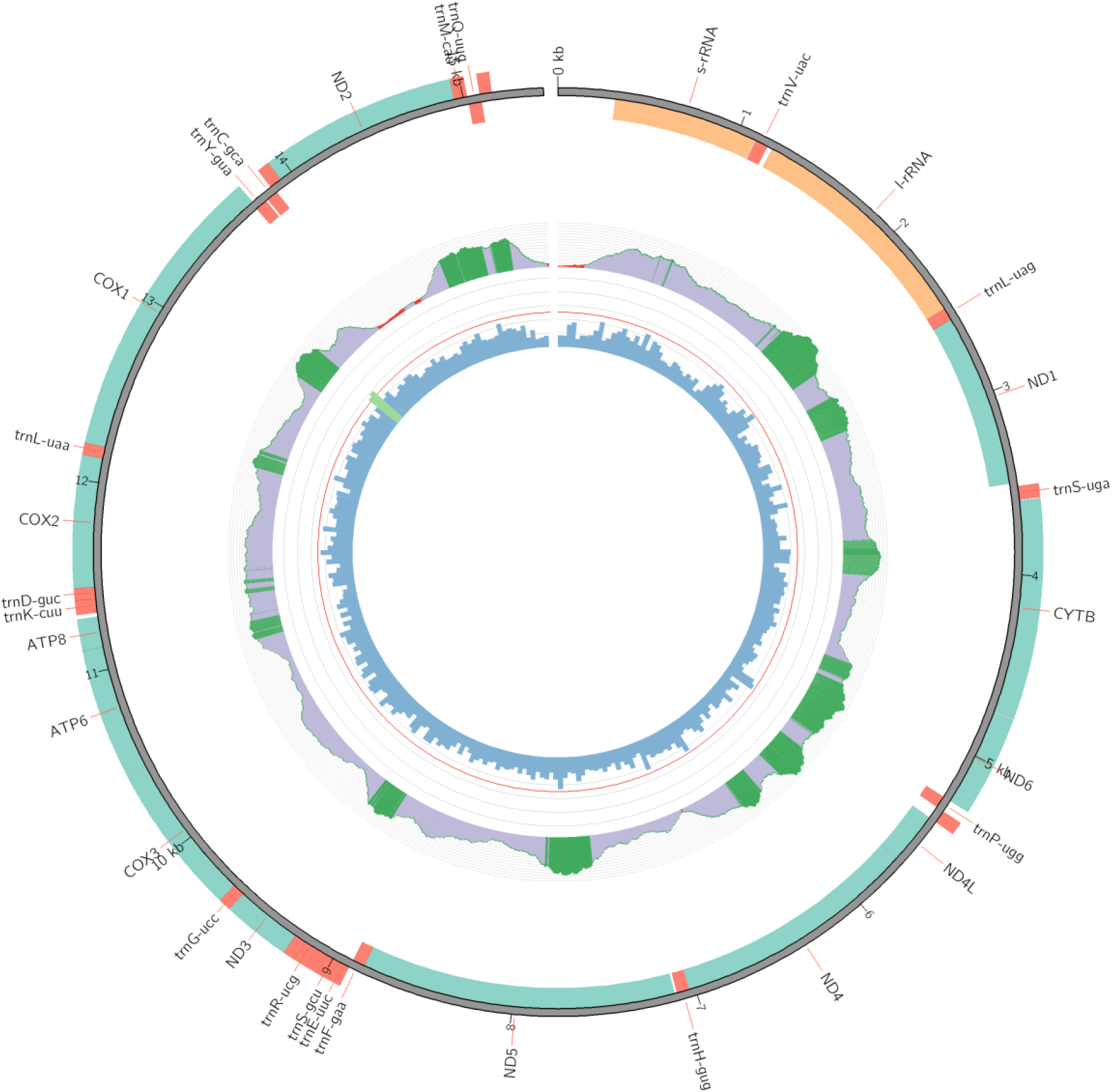
Demonstration of mitogenome visualization using MitoZ.

### Comparison between MitoZ and NOVOPlasty

NOVOPlasty successfully recovered 570 (76.00%) PCGs and rRNA genes of full length, partially assembled 30 (4.00%) genes and failed to assemble 150 (20.00%) genes (Fig. 5 (i)). A total of 133 (17.73%) NOVOPlasty-failed genes were successfully assembled by MitoZ. The gene similarities between NOVOPlasty and Sanger mitogenomes were in concert with that of MitoZ, indicating these genes of low similarities were probably attributed to intra-species genetic variation (Fig. 5 (ii)).

**Fig. 5.**
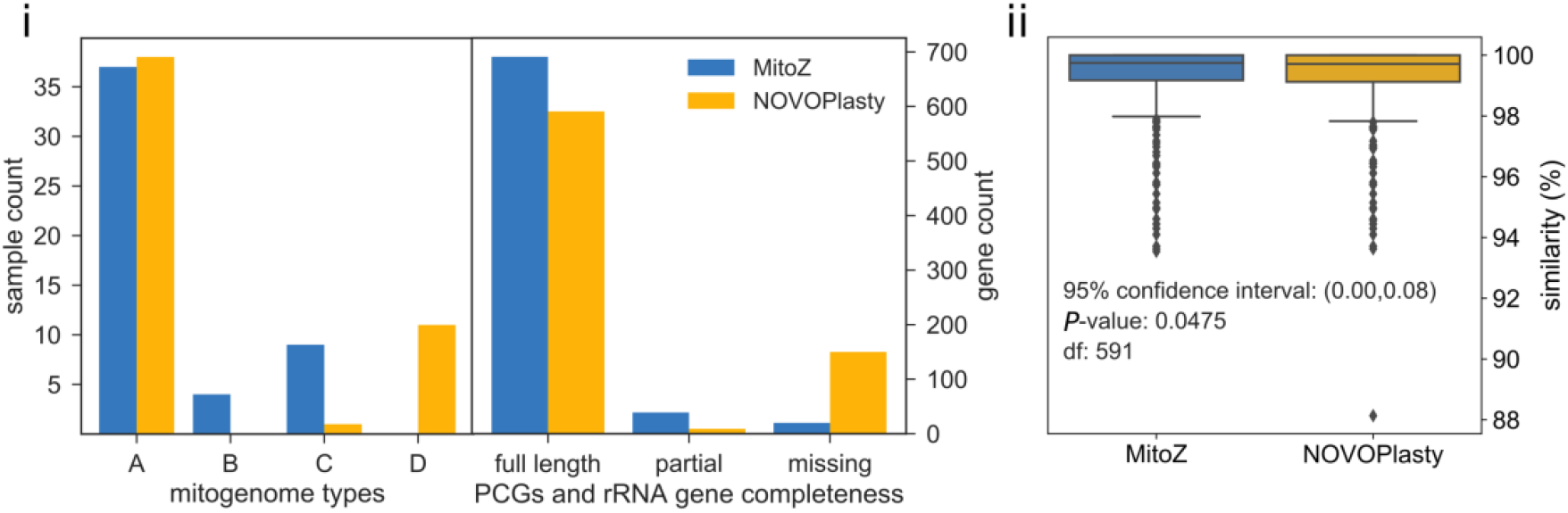
Performance comparisons between MitoZ and NOVOPlasty. (i) Left: mitogenome types, see Fig. 2(i) and Fig. 2(ii) for the categories of mitogenome types, while type D indicates the total number of PCG and rRNA genes recovered was less than eight. Right: gene (PCGs and rRNA genes) completeness distribution, see Fig. 2(iii) for the meanings of “full length”, “partial” and “missing”. (ii) Diagram of gene similarities to the Sanger genes. Genes that recovered by both MitoZ and NOVOPlasty (n=592) were included in the analysis, and paired samples t-test was used.

### Factors influencing mitogenome assembly

The portion of mitochondrial DNA varies in different samples. Our results showed that the assemblies of type A and B mitogenomes tended to possess higher MDR ratios than the assemblies of type C and D (Fig. S2 (i) and (ii)). However, no significant correlation was detected between mitogenome assembly qualities and factors of both A+T content and heterozygosity ratio (Fig. S2 (iii) and (iv)).

## DISCUSSION

The development of HTS technique and low sequencing cost greatly facilitates mitogenome sequencing which used to require a tedious process of long range PCR followed by primer walking (56). Nowadays, we are able to obtain mitogenomes from shotgun reads with higher efficiency and lower cost. At the same time, however, the large volume of data challenges our ability to efficiently analyze the exponentially growing dataset. MitoZ, a versatile mitogenome toolkit, aims to achieve mitogenome assemblies together with annotation results from whole genome shotgun reads. It is by far the easiest method to deliver human-readable outcomes for mitogenome studies – require no special pretreatment in either DNA extraction or nucleotide sequencing, and it combines the key bioinformatics steps – clean data filtering, *de novo* assembly, annotation and visualization, thus provides users an ‘one-step’ solution from raw data to publishable outcomes and will accelerate the accumulation of mitogenomes. In addition, MitoZ conducts assembly without the aid of reference sequences from closely-related species, which can be a crucial feature when the mitochondrial genes from closely-related species are unavailable.

The boundaries of PCGs, especially the stop codons, of many mitochondrial PCGs are not precisely determined in Genbank. Aside from tBlastn and Genewise, MitoZ developed an in-house Python script to further determine the start and stop codons around the boundary of each gene and in most cases was able to precisely locate the start and stop codons (Table S7 and Table S8). In addition, the current annotation module, especially PCGs annotation, works well mainly for arthropods and mammals and needs further improvement to support more domains of life.

In summary, MitoZ shows the ability to assemble and annotate mitogenomes efficiently and accurately. With the rapid accumulation of mitogenomes and robust reference databases of specific environments or groups, it will facilitate the developments of several important fields in the foreseeable future, such as phylogenetic inference, quarantine inspection, aquatic and agriculture ecosystems scrutiny.

## Supporting information

## AUTHOR’S CONTRIBUTION

S.L. and G.M. designed this study. Most of the coding work was conducted by G.M., with minor contributions from S.L., Y.L. and C.Y. S.L. and G.M. drafted the manuscript. All the authors contributed to the manuscript revise.

## ACKNOWLEDGEMENT

We sincerely thank to Chengran Zhou, Min Tang and Xin Zhou for their trying out the MitoZ’s beta version and their positive feedbacks. S.L., G.M. and C.Y. is supported by Free-oriented Project from Shenzhen Government (JCYJ20170817150755701).

## Supplementary files

Fig. S1. Sequencing depth and A+T content around the indel regions.

Fig. S2. Factors influencing mitogenome assembly.

Table S1. Species list used for MitoZ performance testing

Table S2. A+T content of sanger mitogenome and sanger mitochondrial genes, mitochondrial reads ratio and heterozygosity of SRA data

Table S3. Procedures and species list for selecting test species for arthropods

Table S4. Procedures and species list for selecting test species for mammals

Table S5. Species list for building PCGs HMM models

Table S6. Species list for building mitochondrial PCGs annotation database Table S7. MitoZ genes vs. sanger genes of 29 arthropod samples

Table S8. MitoZ genes vs. sanger genes of 21 mammal samples

Table S9. Information of the five false positive genes and samples with low similarity (<97%) genes in MitoZ assembly

Table S10. NOVOPlasty genes vs. sanger genes of 29 arthropod samples

Table S11. NOVOPlasty genes vs. sanger genes of 21 mammal samples

