## Supplementary material for "MitoZ: A toolkit for mitochondrial genome assembly, annotation and visualization": Supplementary Figures.pdf

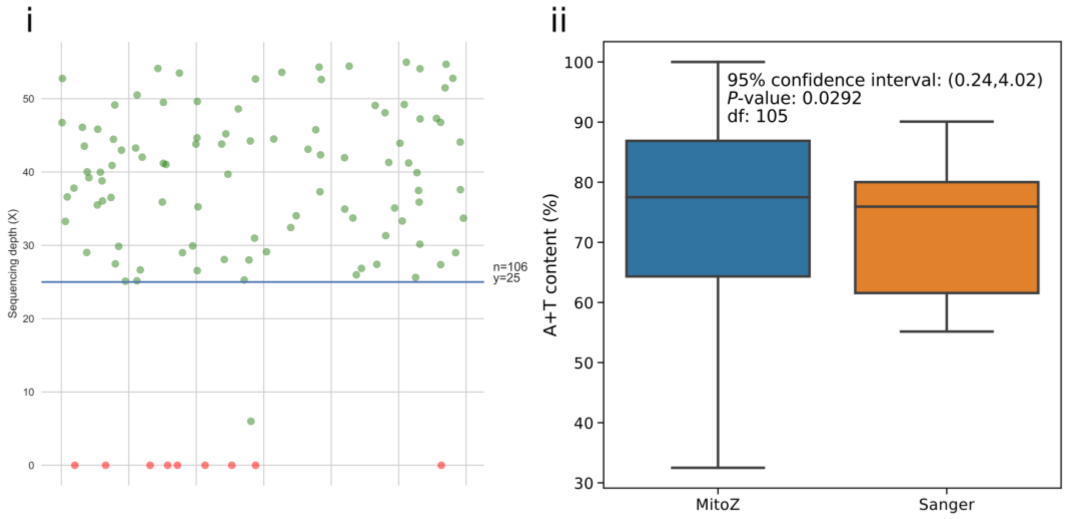


**Fig. S1. Sequencing depth and A+T content around the indel regions.** (i) Depth distribution around the indel regions. We calculated depths of each site around the regions (-20bp, +20bp) and sites of minimum depths were used to represent the depths for each region. Regions of depth ⩾ 40X will be designed a random value between 25 and 55 for better visualization. Red dots indicate regions of sequencing depth of 0X due to 'Ns' in the reference mitogenomes. (ii) A+T content comparison for the indel regions. Paired sample t-test was used, following https://pythonfordatascience.org/paired-samples-t-test-python/.

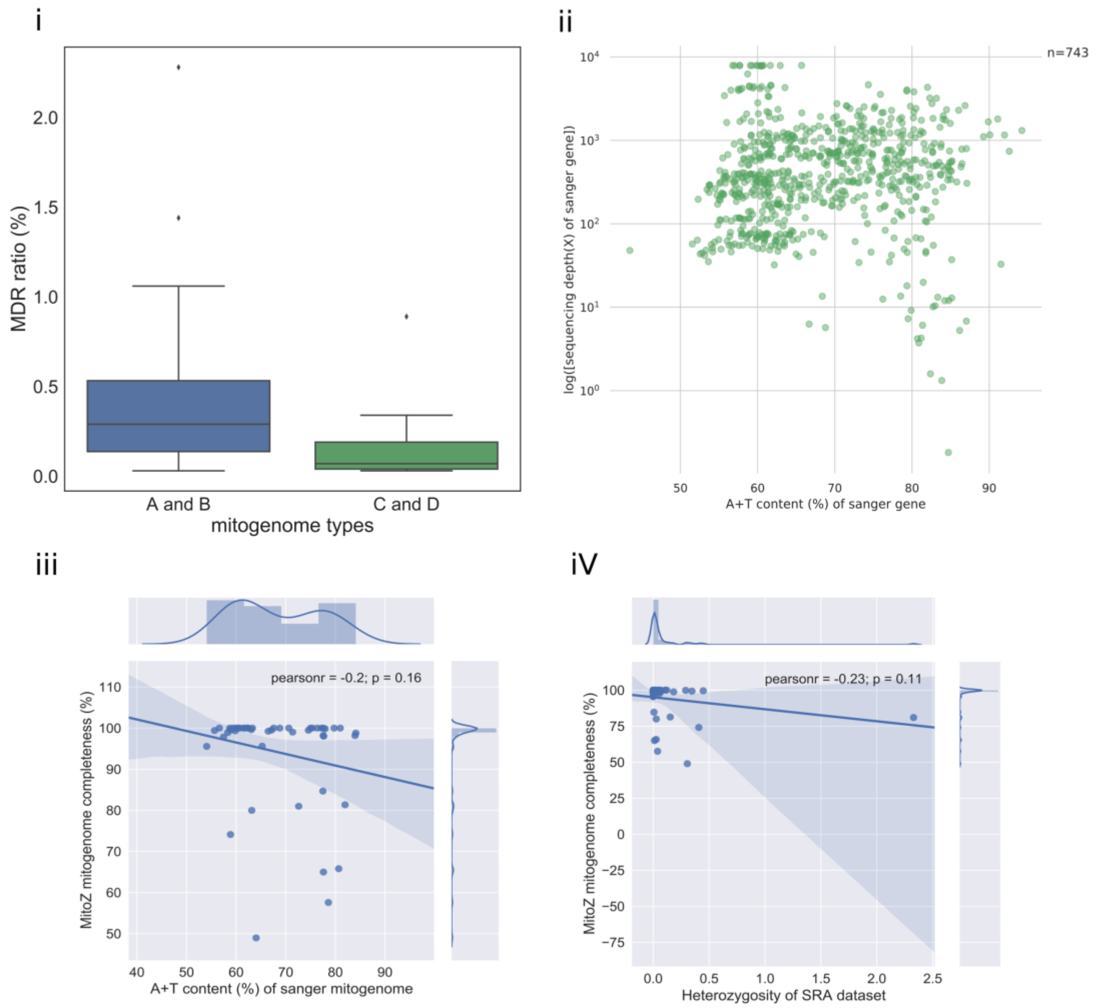


**Fig. S2. Factors influencing mitogenome assembly.** (i) Mitogenomes of types A and type B tend to have higher MDR ratios than mitogenomes of type C and type D. See Fig. 1 for definition of mitogenomes. (ii) Almost all sanger genes have high (⩾5X) sequencing depth, measured by mapping reads to sanger mitongeomes with BWA (version 0.7.12-r1039) with default parameters and using the “samtools depth” command in SAMtools (Li et al. 2009) (version 1.7) to retrieve the sequencing depth. There were six sanger genes were Ns and one ATP8 gene was not annotated, thus only 743 genes were shown. (iii) Relationship of MitoZ mitogenome completeness and A+T content. (iv) Relationship of MitoZ mitogenome completeness and heterozygosity. "Heterozygosity" was measured using ANGSD (Korneliussen, Albrechtsen & Nielsen 2014).
